# A Unified Single-Cell Atlas of HNSCC: Uncovering HPV and Sex Variability in the Tumor Microenvironment

**DOI:** 10.1101/2025.10.30.685278

**Authors:** Cristina Conde Lopez, Divyasree Marripati, Maria Jose Besso, Mareike Roscher, Rui Han, Wahyu Wijaya Hadiwikarta, Moshe Elkabets, Jochen Hess, Ina Kurth

**Affiliations:** German Cancer Research Center Heidelberg (DKFZ), Radiooncology/Radiobiology, Germany; Department of Otorhinolaryngology, Head and Neck Surgery, Heidelberg University Hospital, Heidelberg, Germany; The Shraga Segal Department of Microbiology, Immunology, and Genetics, Ben-Gurion University of the Negev, Beer-Sheva, Israel; Faculty of Health Sciences, Ben-Gurion University of the Negev, Beer-Sheva, Israel Department of Otorhinolaryngology, Head and Neck Tumors, Heidelberg University Hospital, 69120 Heidelberg, Germany; German Cancer Research Center Heidelberg (DKFZ), Service Unit for Radiopharmaceuticals and Preclinical Studies, Germany; German Cancer Consortium (DKTK), Core Center Heidelberg, 69120 Heidelberg, Germany

## Abstract

Head and neck squamous cell carcinoma (HNSCC) is highly heterogeneous, with variations driven by HPV status and sex. However, existing single-cell RNA sequencing (scRNA-seq) studies are often limited in sample size and lack standardized methodologies, limiting cross-study comparisons. To address this, we integrated scRNA-seq data from 78 patients (274,911 cells) across multiple studies, creating a unified HNSCC atlas that harmonizes annotations and enables robust tumor microenvironment (TME) analyses. Using STACAS for semi-supervised integration and automated annotation tools such as Ikarus and scGate, we improved tumor and immune cell classification. Leveraging our atlas, we identified HPV- and sex-specific shifts in immune and stromal composition, with HPV+ tumors enriched in adaptive immune cells and HPV− tumors showing more stromal and myeloid populations. Preliminary sex-stratified analyses suggested distinct microenvironmental patterns, warranting further investigation. This publicly available atlas provides a comprehensive framework for reproducibly studying HNSCC biology, improving patient stratification, and may help informing personalized therapies.

## Introduction

Head and neck squamous cell carcinoma (HNSCC) accounts for over 90% of head and neck cancers (HNCs), with human papillomavirus (HPV) infection as a major etiological risk factor, particularly in oropharyngeal cancers^1–3^. HNSCC also exhibits significant sex differences, with a higher incidence in males, traditionally linked to tobacco and alcohol use, though recent evidence suggests additional biological and molecular contributors^4,5^. Despite advances in treatment, the five-year overall survival (OS) remains below 50%, and relapse rates exceed 50%, indicating the need for improved therapeutic strategies^1^. Managing HNSCC is particularly challenging due to high intra- and inter-tumor heterogeneity, yet current treatment approaches remain largely uniform, highlighting the necessity to improve outcomes through better patient stratification^6,7^.

For the past decade, transcriptomics has largely relied on bulk RNA sequencing, which captures aggregate gene expression but obscures cellular heterogeneity. While collaborative initiatives like The Cancer Genome Atlas (TCGA) have identified tumor subtypes and molecular signatures, bulk RNA sequencing cannot resolve the distinct contributions of different cell types, limiting its ability to reveal key therapeutic targets^8–13^. Single-cell RNA sequencing (scRNA-seq) has transformed cancer research by enabling high-resolution analysis of the tumor microenvironment (TME) ^14^. This approach uncovers interactions between tumor, immune, and stromal cells, which are critical for cancer progression and treatment response^15^. However, inconsistencies in sample sizes, annotation standards, and methodologies across studies complicate the unified understanding of HNSCC through scRNA-seq. For example, variations in chromosomal aberrations, HPV gene expression, and macrophage polarity have been identified across studies, yet differences in focus and specifically cellular annotations have resulted in fragmented knowledge, emphasizing the need for a standardized framework^2,16^.

HNSCC research faces several challenges, including high intra- and inter-patient TME heterogeneity driven largely by tumor cells and the stromal component. Another major hurdle is the inconsistency in tumor cell annotation, as classification criteria vary based on gene expression and copy number variation (CNV) patterns. The lack of standardization across studies affects data comparability and impedes the progress in targeted therapy development. Machine learning-based approaches offer potential solutions for improving tumor cell annotation consistency, while the integration of multi-study datasets helps address limitations of small sample sizes^14,17,18^. Additionally, the dynamic nature of cells presents a persistent problem, as scRNA-seq captures only snapshots of cellular states, complicating comparability across studies^19^. Advances in AI-driven annotation tools and community-led initiatives, such as “Single Cell Best Practices” and “scRNA-tools,” aim to establish standardized workflows and improve data harmonization^20,21^. Resource-intensive single celltechniques remain a challenge, but public datasets and computational strategies like pseudobulking offer cost-effective alternatives to enhance uniformed data reusability.

In this study, we present our newly developed pipeline for integrating multiple single cell datasets to create a comprehensive atlas of HNSCC, with the aim of overcoming the limitations of isolated studies. By consolidating data from various independent studies into a unified “universal atlas”, we achieve a holistic view of the TME. By incorporating key variables such as HPV status and sex, the atlas provides a powerful tool for unraveling the underlying biological and molecular mechanisms that drive etiological factors and their impact on tumor behavior and patient outcomes. This atlas provides a high-resolution map of cellular diversity and functional interactions within the TME, bridging gaps in existing research and enhancing our understanding of cancer progression and treatment resistance. The pipeline and atlas are publicly available (https://github.com/DKFZ-E220/HNSCatlas) as a resource for the research community to facilitate further investigation and discovery.

## Results

### Generation and Validation of the HNSCC Atlas

To create a comprehensive HNSCC atlas, a systematic literature search was conducted, and four publicly available scRNA-seq datasets were selected (GSE234933, GSE182227, GSE164690, and GSE181919), they were chosen for their detailed clinical annotations, including sex, HPV status, and sample origin (normal tissue, primary tumor, or metastasis). This approach integrated data from 78 patients with 274,911 cells as data points significantly expanding beyond the typical range of 11–20 patients in individual HNSCC datasets (Suppl. Table 1). Important data curation steps, such as removing low-quality data points and harmonizing variables, ensured a consistent data structure across studies, providing a strong foundation for integration. These initial steps of quality control, normalization, and scaling were performed using the Seurat pipeline. A schematic view of the atlas creation process can be found in Fig. 1.

**Figure 1.**
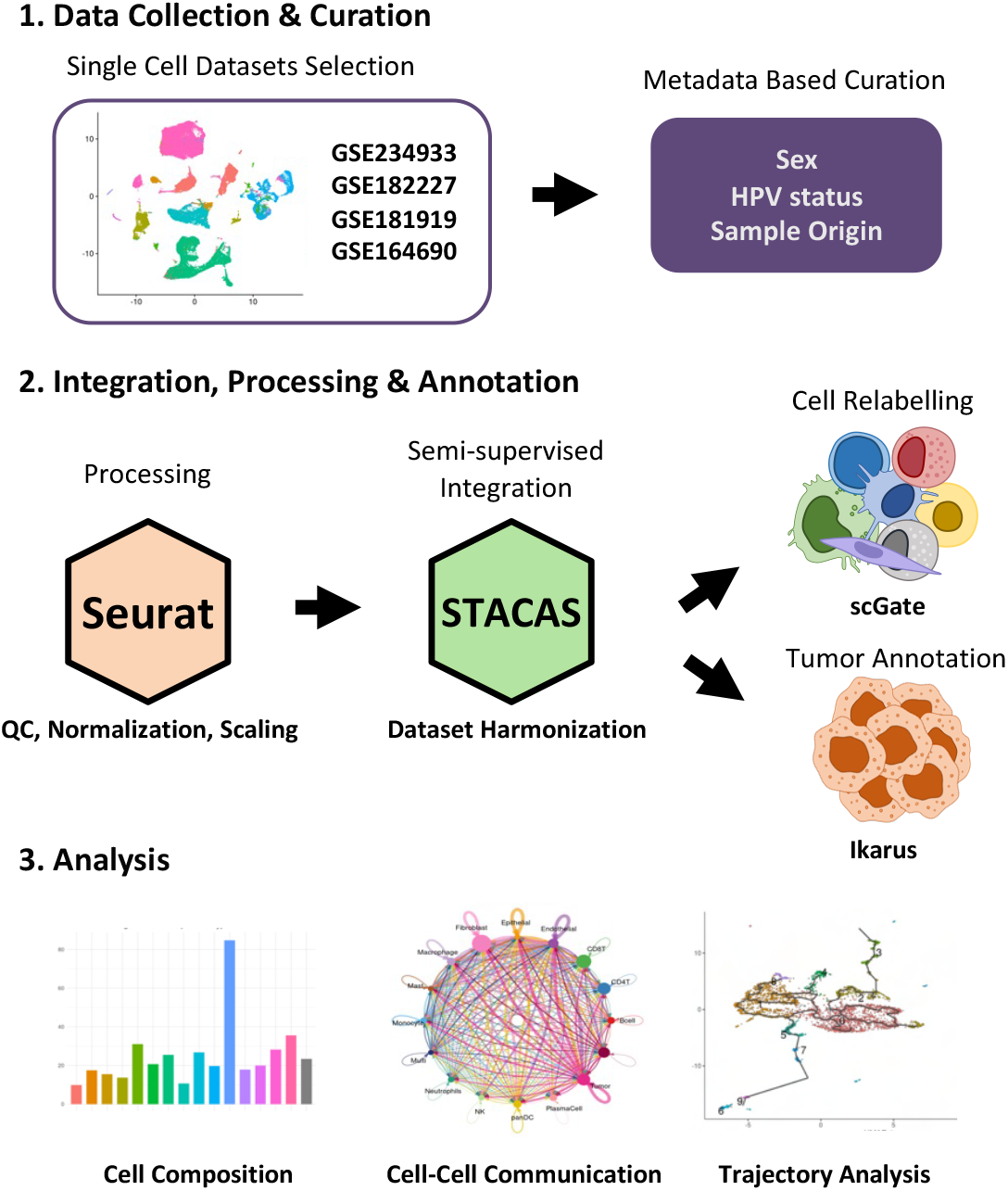
Workflow for creating the HNSCC atlas. The workflow involves three main stages. First, single-cell RNA-seq datasets are collected and curated based on key metadata, including sex, HPV status, and sample origin. Next, data processing is performed with Seurat for quality control, normalization, and scaling, followed by dataset harmonization using STACAS for semi-supervised integration. Cell relabeling is refined with scGate, and tumor cells are specifically annotated using Ikarus. Finally, the harmonized atlas allows comprehensive downstream analyses, for example cell composition profiling, cell-cell communication inference, and trajectory analysis.

The harmonization process significantly improved data compatibility across the HNSCC single cell datasets, as shown in the Uniform Manifold Approximation and Projection (UMAP) plots (Fig. 2 A, B). The integration quality was quantitatively validated using established metrics such as LISI and ASW, confirming superior performance of the STACAS pipeline compared to alternative methods (Supplementary Fig. 1). This refinement led to a more cohesive clustering of cells, with increased density within each cell type and improved separation between distinct populations. While the number of annotated cell types decreased from 24 to 16, this reduction reflects the resolution of inconsistencies in initial classifications, leading to a more biologically coherent representation of the TME. The harmonization process improved consistency across studies by standardizing original annotations to a common framework (Fig. 2A), while scGate and Ikarus enabled further refinement of cell identities (Fig. 2B). This was particularly relevant for tumor cells, which were underrepresented in the original labels. Application of Ikarus, a model-based classifier, expanded the annotated tumor cell population from 2,300 to nearly 50,000 cells, most of which clustered within epithelial regions of the UMAP. These improvements are also reflected in the mosaic plot (Fig. 2C), where a clear shift from epithelial to tumor identity is observed in the relabeled dataset.

**Figure 2.**
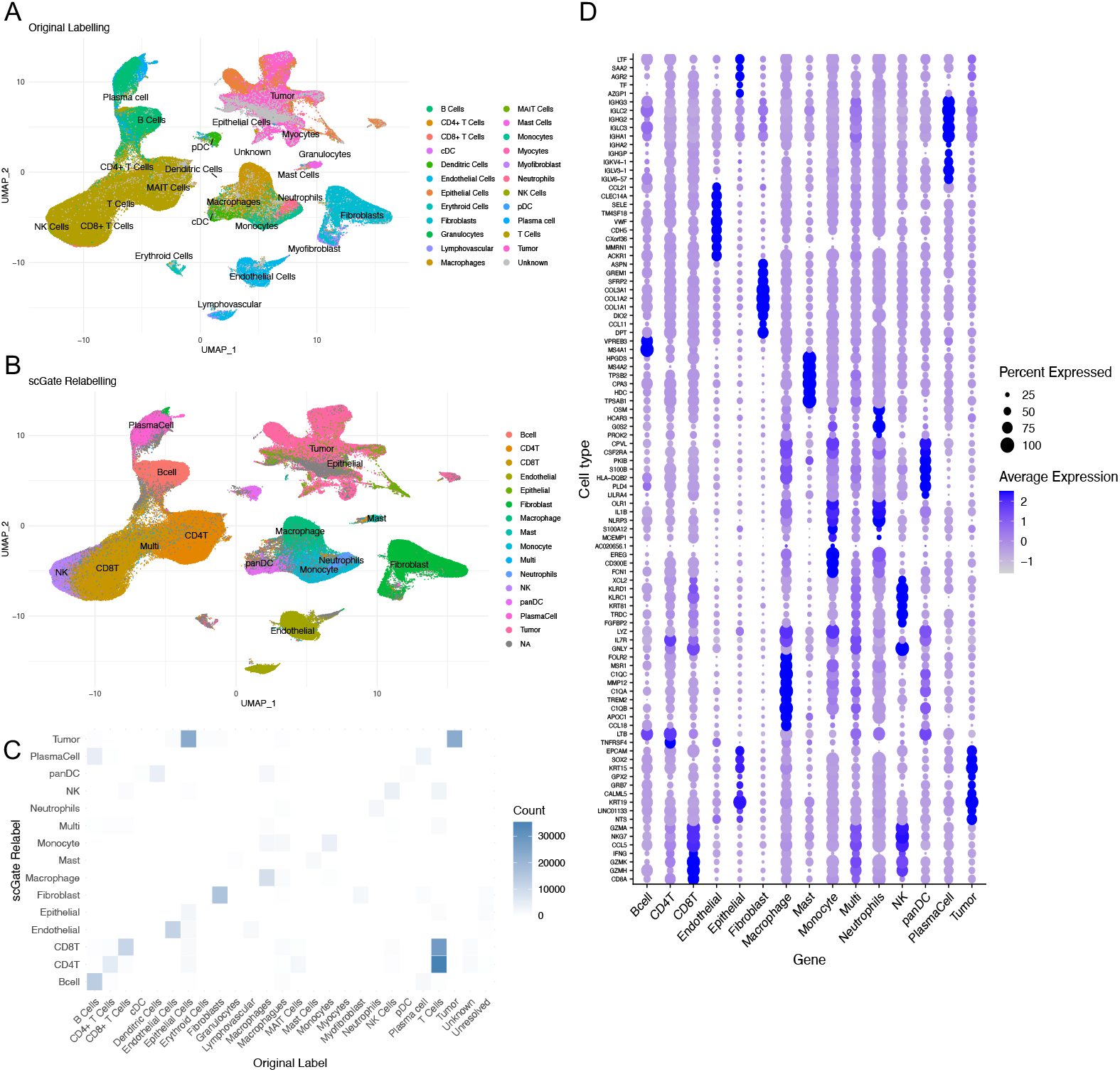
Integration and cell relabeling in the HNSCC atlas using scGate. **(A)** UMAP visualization of the original cell type labels across the integrated single-cell RNA-seq datasets. **(B)** UMAP showing the updated cell annotations following scGate-based relabeling, with an additional tumor cell population annotated using the Ikarus classifier. This relabeling enhanced the resolution of tumor and non-tumor cells, particularly by reclassifying epithelial cells as tumor cells where appropriate. **(C)** Mosaic plot comparing the distribution of original and scGate-relabelled cell types. The plot illustrates the reassignment of cells, especially the transition of epithelial cells to the tumor cell category. **(D)** Dot plot showing top differentially expressed genes across annotated cell types. This analysis highlights the most enriched markers in each major cell population.

### Automated Annotation of Cell Types

TME cell types were annotated using automated gating models from scGate, which applies signature-based scoring to assign cells to predefined populations. This relabeling process refined the original annotations by resolving inconsistencies and enhancing cell-type resolution across samples. The UMAP plots (Fig. 2A, B) illustrate the improvement in label resolution across the atlas, particularly for challenging populations such as T cell subtypes and fibroblasts. To evaluate the global effect of the relabeling process, the original and updated annotations were compared using a mosaic plot (Fig. 2C), highlighting key transitions such as the separation of overlapping plasma and B cells. The T cell compartment, originally annotated as a single cluster, is now split into CD4 and CD8 subsets, reflecting increased resolution and consistency in immune classification across studies. To quantitatively assess the extent and consistency of these refinements, the Adjusted Rand Index (ARI = 0.338) and Normalized Mutual Information (NMI = 0.516) between the original and relabeled annotations were computed. These values indicate moderate agreement, consistent with biologically meaningful restructuring rather than simple label preservation and support the improved coherence of the final annotations. To validate the final annotations, differential expression analysis was performed across all major cell types, identifying top markers such as CD8A, GZMH, and GZMK in CD8^+^ T cells, LTB and TNFRSF4 in CD4^+^ T cells, and ASPN, GREM1, SFRP2, COL3A1, and COL1A2 in fibroblasts (Fig. 2D). Signature scoring further confirmed the selective enrichment of these gene sets within the expected populations (Supplementary Fig. 2), supporting the biological specificity and robustness of the automated annotation.

### inferCNV Analysis for Tumor Cell Reclassification

Since tumor cells typically exhibit higher chromosomal instability compared to normal epithelial cells, chromosomal copy number variation (CNV) analysis can serve as an independent validation of transcription-based classification methods. Although reclassified tumor cells showed strong tumor-like expression signatures (Supplementary Fig. 2) and were confidently identified by the Ikarus algorithm, we further validated their classification using CNV analysis. We analyzed three groups: (1) originally annotated tumor cells, (2) epithelial cells, and (3) cells reclassified from epithelial to tumor using the Ikarus algorithm. The analysis confirmed that tumor cells identified using the Ikarus algorithm displayed a higher level of chromosomal instability compared to normal tissue epithelial cells ( Fig. 3). This approach validated the utility of the Ikarus method, as it enables the identification of cells with significant CNV based on gene expression.

**Figure 3.**
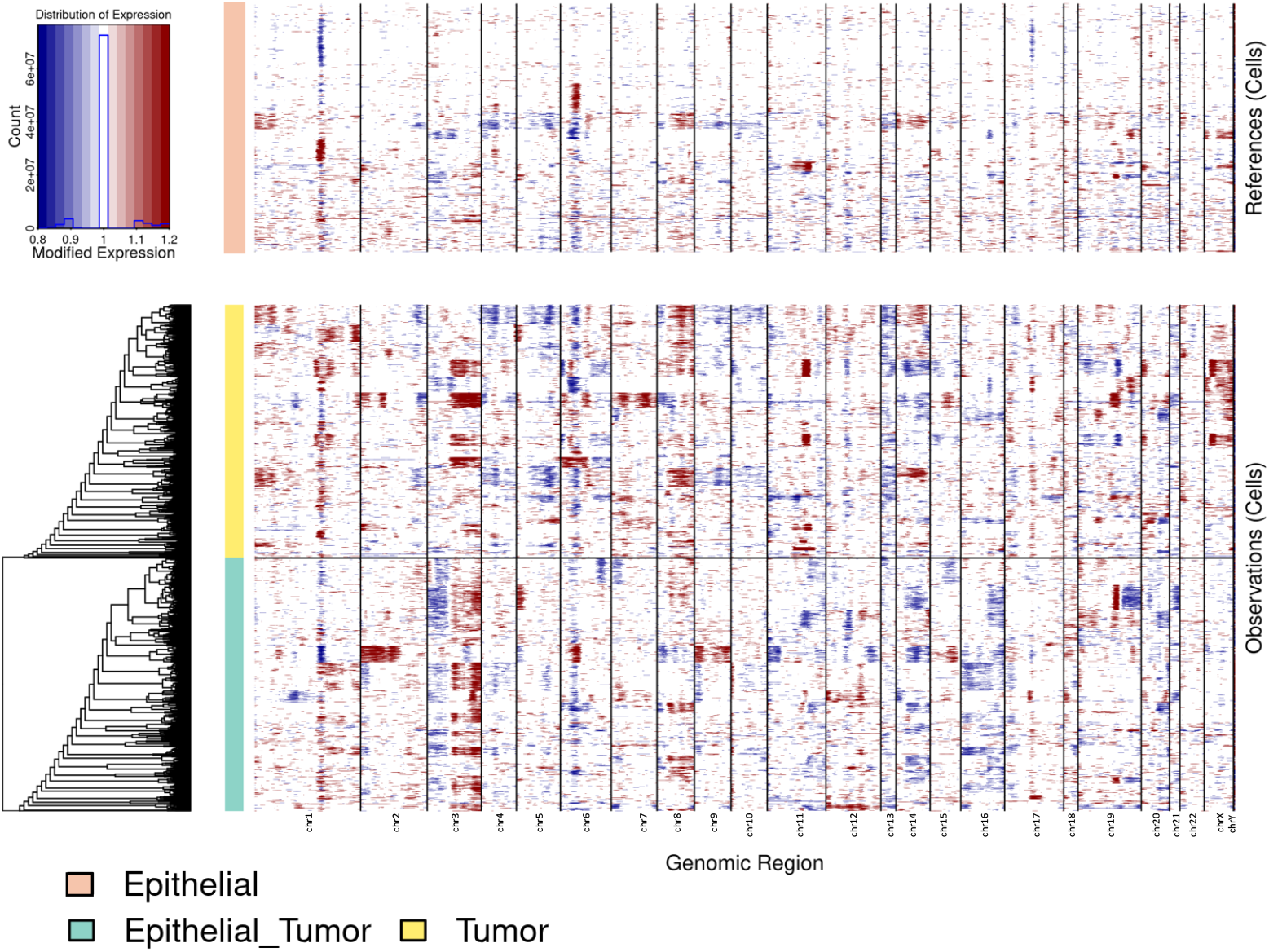
Validation of tumor cell reclassification using inferCNV analysis. The copy number variation (CNV) analysis was performed on epithelial cells, originally annotated tumor cells, and cells reclassified from epithelial to tumor. Heatmaps show chromosomal instability across genomic regions, with red and blue representing gains and losses, respectively. Cells reclassified as tumor using the Ikarus algorithm displayed a higher level of chromosomal instability compared to epithelial cells, validating their tumor identity. This approach demonstrates the utility of Ikarus for identifying tumor cells based on gene expression-derived CNVs.

### Refinement of Immune Compartment Annotations

Using gating models from scGate, the classification of immune cells was refined and deepened. Predefined subtype signatures from frameworks like ProjectTILS were used to label key types such as CD4 and CD8 T cells and dendritic cells, ensuring consistent and reliable annotations across our dataset. To further validate the annotation of T cell subtypes, differential expression analysis was performed to identify top markers for each cluster, which showed strong concordance with expected gene expression patterns (Supplementary Fig. 3). A complete list of cell subtype abbreviations used in this study is provided in Supplementary Table 6.

### Deep TME Analysis Across Datasets Using the Unified HNSCC atlas

Next, the unified atlas was used to conduct an in-depth TME analysis of the 78 HNSCC patients overcoming previous limitations imposed by the small sample sizes of individual datasets and inconsistencies in cell annotation. The cohort included 46 HPV- and 32 HPV+ tumors, with a strong male predominance (62 males, 16 females), which limited the statistical power of some comparisons.

To assess differences in cell type composition between HPV+ and HPV− tumors, per-patient proportions were computed by dividing the number of cells of each type by the total number of cells per patient (Fig. 4A), capturing interindividual variability in tumor composition. To complement this, cell type proportions were scaled to a fixed total (Fig. 4B), generating simulated counts that provided a more standardized measure of relative abundance. While simulated counts reduced interpatient variation, patient-level proportions offered valuable context for individual heterogeneity. Both approaches revealed consistent trends with HPV+ tumors showing enrichment in adaptive immune cells, including B cells, plasma cells, and CD4/CD8 T cells, suggestive of a more active immune microenvironment, whereas HPV-tumors showed elevated levels of innate and stromal populations such as macrophages, monocytes, fibroblasts, and neutrophils, aligning with a more suppressive TME^22^. NK cells were also more abundant in HPV+ tumors. To further explore TME heterogeneity, cell composition was stratified by both HPV status and patient sex (Suppl. Fig. 4, Suppl. Table 2–6), revealing sex-specific differences in fibroblast and tumor cell abundance as well as varying levels of exhausted versus cytotoxic CD8 T cells, particularly in HPV+ tumors.

**Figure 4.**
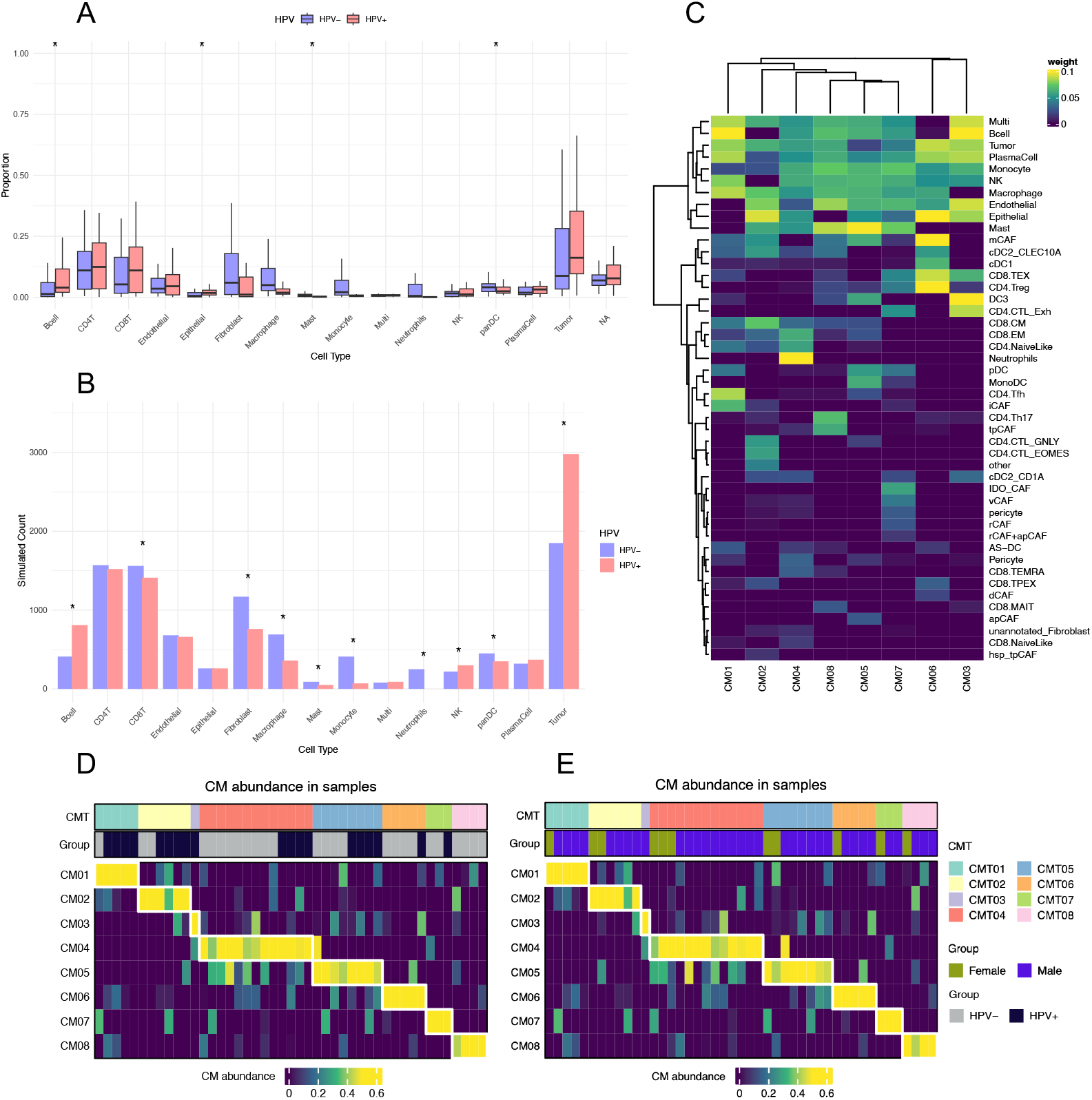
HPV-specific cell type distributions and higher-order coordination patterns in the HNSCC TME. **(A)** Boxplots presenting per-patient cell type proportions in HPV-positive (HPV+) and HPV-negative (HPV-) tumors. Statistical comparisons were performed using the Wilcoxon test, and significance is indicated as p < 0.05 (*). **(B)** Barplots showing projected cell counts based on proportions for each cell type in HPV+ and HPV-tumors. Statistical comparisons were performed using the Fisher’s exact test, and significance is indicated as p < 0.05 (*). **(C)** Heatmap of CoVarNet-derived module weights across cell types, showing the contribution of each cell type to the eight identified co-variation modules (CM01–CM08). **(D)** Module abundance per patient stratified by HPV status. **(E)** Module abundance per patient stratified by sex.

To investigate coordinated shifts in the TME, CoVarNet was applied, a computational framework that identifies cellular co-variation modules (CMs) by analyzing co-fluctuations in cell type abundances across tumors. This analysis yielded eight distinct co-variation modules (CM01– CM08), each composed of cell types that tend to increase or decrease together across patient samples (Fig. 4C). For example, CM01 was enriched for CD4 Tfh cells, B cells, plasma cells, macrophages, and notably iCAFs, indicative of a humoral and immunoregulatory microenvironment. CM04 was dominated by neutrophils and CD8 EM cells, pointing to a more inflammatory composition. CM06, on the other hand, integrated CD4 Tregs, mCAFs, and CD8 TEX, potentially reflecting an exhausted or immunosuppressive microenvironmental niche.

The distribution of module abundance across tumors showed that CM01, CM02, and CM03 were more prominent in HPV+ tumors, whereas CM04, CM06, CM07, and CM08 were more abundant in HPV-tumors, suggesting that HPV-tumors may harbor more immunosuppressive or inflammatory TMEs, while HPV+ tumors are characterized by more coordinated lymphoid and stromal ecosystems (Fig. 4D). When stratified by sex (Fig. 4C), no clear pattern emerged; all modules were represented in both male and female tumors, with no strong enrichment apart from CM03, which was present in only a single male sample and is therefore not interpretable.

Building on the coordinated ecosystem patterns identified above, the analysis next focused on male patients to explore differences in cell composition and immune activity between HPV+ and HPV-tumors. We restricted this analysis to males (n = 62; 32 HPV− and 30 HPV+) due to the limited number of female samples, which did not allow for a balanced comparison. Cell type proportions revealed significant differences between the two groups (Fig. 5A): HPV+ tumors showed increased abundance of tumor cells and B cells, while HPV− tumors were enriched for monocytes, macrophages, mast cells, and fibroblasts, pointing toward a more inflammatory and stromal-rich microenvironment in the absence of viral infection. To further dissect immune cell states, we evaluated pathway-level activity in CD4 and CD8 T cells using UCell scoring. Although scores varied across patients (Fig. 5B), both the cytotoxic CD8 T cell and Treg signatures were significantly elevated in HPV-tumors (Fig. 5C), suggesting a coexistence of cytotoxic and immunosuppressive programs within the same tumor context, a pattern that contrasts with previous reports of enhanced cytotoxicity in HPV+ tumors. This discrepancy may reflect differences in cohort composition, sample processing, or tumor-intrinsic heterogeneity, and warrants further investigation.

**Figure 5.**
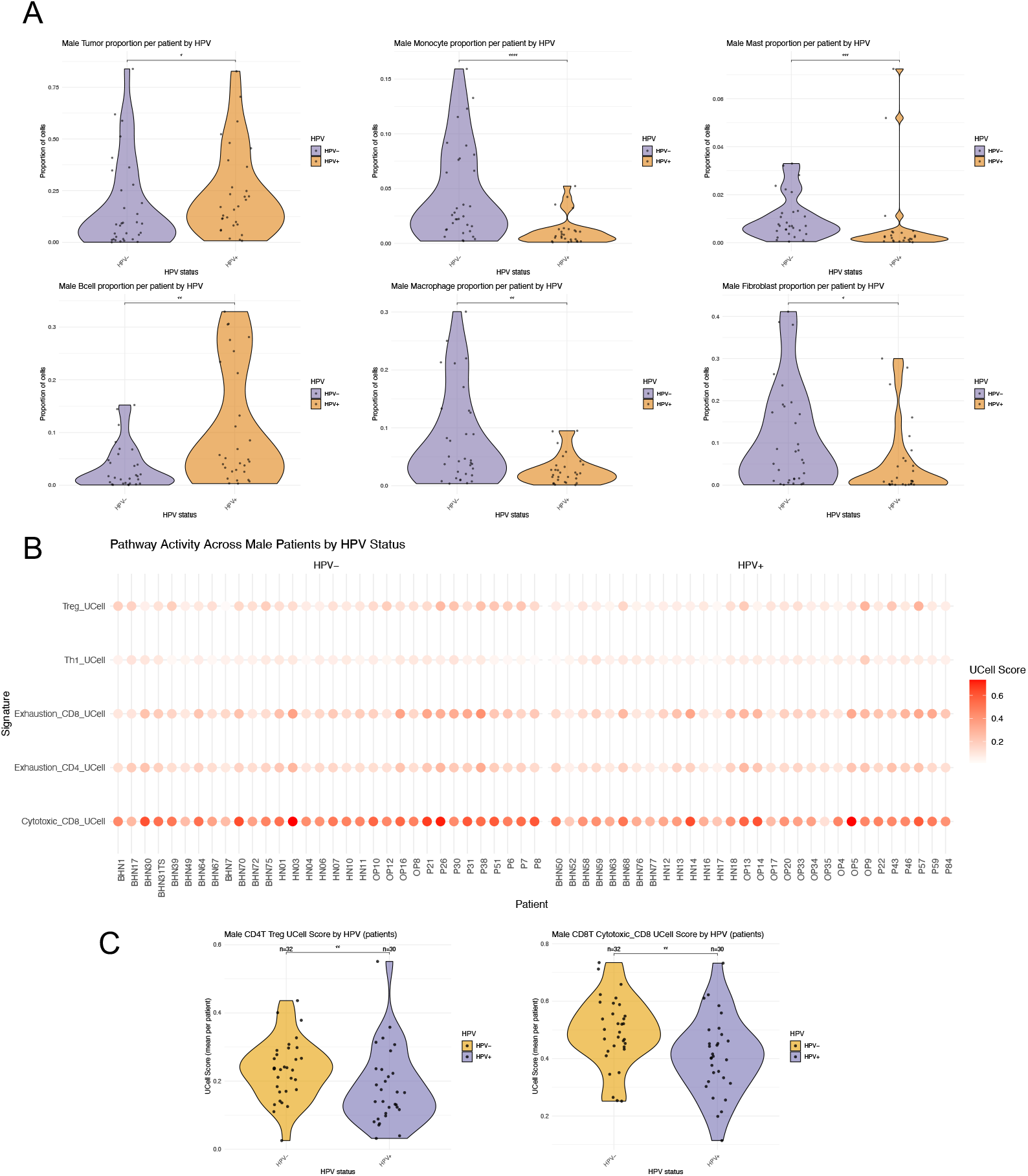
HPV status shapes immune cell composition and functional states in male HNSCC tumors. **(A)** Violin plots showing the relative proportions of major immune cell types across HPV+ and HPV− male patients. Significant differences were determined by Wilcoxon rank-sum test. **(B)** Dotplot displaying per-patient UCell scores for selected CD4^+^ and CD8^+^ T cell gene signatures (Treg, Th1, exhaustion, and cytotoxicity), stratified by HPV status. **(C)** Violin plots of Treg and cytotoxic CD8 T cell signature scores, which showed statistically significant differences between HPV+ and HPV groups (Wilcoxon test).

## Discussion and Outlook

The creation of a comprehensive single-cell atlas marks a significant advancement in the study of HNSCC. By integrating data from 78 patients and 274,911 cells across diverse clinical backgrounds, we have created a high-resolution map of the TME. This atlas represents an important step toward addressing dataset heterogeneity, offering a standardized framework for systematically exploring cellular composition and function across cohorts. This approach not only enhances the resolution of cellular phenotypes and facilitates the exploration of inter dataset variability but also allows for the robust validation of findings across diverse patient samples and experimental conditions.

Compared to previous efforts in integrating single-cell data from HNSCC samples, such as the work by Dai et al., the approach stands out for incorporating an integration strategy and refined annotation methodologies that build upon existing approaches^23^. While Dait et al. integrated five datasets using the standard Seurat pipeline with manual annotation based on cluster markers, we expanded upon this by incorporating STACAS for semi-supervised integration, leveraging existing labels to guide the harmonization process. In addition, we implemented automated annotation pipelines, including the Ikarus algorithm and scGate to ensure consistency and reproducibility across datasets. The reannotation of some epithelial cells as malignant was validated using inferCNV, which confirmed higher CNV counts, indicative of their tumorigenic nature. This novel workflow enhances the identification of more detailed cellular interactions and phenotypes, significantly improving the resolution at which TME dynamics can be studied.

We have generated a comprehensive single-cell atlas that integrates and harmonizes diverse scRNA-seq datasets from multiple HNSCC cohorts. This unified resource enables a holistic and stratified analysis of the TME, overcoming prior limitations imposed by small sample sizes and inconsistent cell annotations. By incorporating key clinical variables such as HPV status and sex, our atlas allows for a detailed dissection of cellular compositions and immune states, which may help inform future studies on therapy response. The in-depth analysis of 78 tumors (46 HPV−, 32 HPV+), with a predominant representation of male patients (62 males, 16 females), points toward distinct differences in TME composition between HPV groups. HPV+ tumors tended to be enriched in adaptive immune components such as B cells and T cells, whereas HPV-tumors exhibited higher levels of stromal and innate immune populations, including macrophages, monocytes, fibroblasts, and mast cells. These patterns suggest that HPV-tumors harbor a more inflammatory and stromal-rich microenvironment, potentially associated with immune suppression. Our observations are further supported by recent studies suggesting a more robust immune activation in HPV+ cases^24^. To capture higher-order relationships, CoVarNet was applied and allowed for the identification of distinct cellular co-variation modules reflecting coordinated immune and stromal ecosystems differentially abundant in HPV+ and HPV-tumors.

To further explore TME differences, we focused on the male subgroup. This analysis confirmed marked shifts in overall cell type composition, with HPV+ tumors showing increased abundance of tumor cells and B cells, whereas HPV-tumors were enriched for monocytes, macrophages, fibroblasts, and mast cells, patterns indicative of a more inflammatory and stromal-rich TME in HPV-tumors. To further dissect immune function, we examined pathway-level activity in CD4^+^ and CD8^+^ T cells using UCell scoring. Interestingly, both cytotoxic CD8^+^ T cell and Treg signatures were significantly elevated in HPV-tumors. This finding is unexpected, as previous studies have typically reported stronger CD8-mediated cytotoxic responses in HPV^+^ tumors, likely driven by viral antigenicity^25^. The discrepancy may stem from cohort-specific differences or sample handling, and shows the importance of stratified and context-aware analyses. Although the limited number of female patients did not allow for robust statistical comparisons, preliminary trends suggest that HPV-females may harbor a distinct TME, potentially shaped by sex-specific immune regulation. These observations, together with prior studies highlighting sex differences in TME composition and immune behaviour, point to the need for future studies with balanced cohorts to explore whether female patients could benefit from tailored therapeutic approaches^26,27^.

Despite the high resolution of the single-cell atlas, scRNA-seq has inherent technical limitations, including dropouts and sampling bias, which may obscure certain biological signals. Additionally, the high cost and resource-intensive nature of single cell technologies can limit accessibility, potentially introducing biases in sample selection^28^. On the computational side, analyzing large single cell datasets requires substantial storage, computational power and continuously evolving bioinformatics tools to manage the growing complexity of data. Furthermore, study design itself can introduce biases, as seen in this atlas, where the proportion of male and female patients is unbalanced, potentially affecting the interpretation of sex-related differences. This limitation, together with the evolving landscape of available datasets, highlights the importance of ongoing updates. While thus atlas integrates the most comprehensive single cell data available at the time of analysis, recent studies, such as those by Xiong et al. (2024) and Kim et al. (2025), have since added valuable transcriptomic information^29,30^. However, due to the absence of key clinical annotations such as HPV status and the lack of detectable HPV transcripts, they were not suitable for inclusion in our stratified framework. However, the open and modular pipeline allows for the future incorporation of such datasets in complementary analyses. These challenges and opportunities highlight the need for continued refinement of both experimental design and analytical strategies to fully leverage the potential of single-cell data in understanding tumor biology.

One promising avenue to overcome these limitations is the integration of multi-omics data, incorporating genomics, transcriptomics, proteomics, and metabolomics to provide a more comprehensive understanding of the TME. Combining scRNA-seq with spatial transcriptomics or epigenetic profiling, for instance, we could enhance the resolution of tumor-immune interactions and identify regulatory mechanisms shaping immune responses in HNSCC^31^. Additionally, longitudinal studies tracking tumor evolution over time would provide valuable insights into tumor heterogeneity, adaptation to treatment and immune evasion. Addressing these challenges while maintaining a robust integrative approach, such as our unified annotation atlas, will be essential for improving the development of effective, personalized therapies in HNSCC.

In summary, this study highlights the value of large-scale data integration in advancing the understanding of the TME in HNSCC. By creating a unified single-cell atlas, we synthesized data from diverse patient transcriptomic profiles, overcoming the inconsistencies that often hinder cross-study comparisons. This integrative approach enhances statistical power by incorporating a larger number of samples, allowing for more refined stratification and robust subgroup analyses. With comprehensive clinical annotations such as sex, HPV status, and environmental exposures (e.g., alcohol or tobacco), the atlas allows a deeper exploration of factors shaping the TME and addresses a broader range of research questions.

## Conclusion

Single-cell transcriptomics has opened up new opportunities to map the TME in significant detail. By integrating data from 78 patients, we created a unified HNSCC atlas that allowed us to explore key factors like sex, HPV status, and immune and stromal composition on a much larger scale than was previously possible. This integration not only overcomes limitations of isolated studies but also sheds light on distinct immune and stromal profiles across these variables, offering insights into tumor biology that were previously unattainable for HNSCC. Our findings reveal clear differences in TME organization not only between HPV+ and HPV− tumors but also across sexes, particularly within the immune and stromal compartments. These observations point toward sex-specific microenvironmental programs that may influence tumor progression and immune activity, highlighting an often-overlooked variable in HNSCC biology.

This atlas is publicly available to ensure its utility as a resource for the research community, facilitating reproducibility and allowing others to explore new hypotheses in HNSCC. Moving forward, this integrative approach emphasizes the importance of harmonized datasets.

## Methods

### Data Collection and Pruning

A comprehensive search was conducted for publicly available HNSCC scRNA-seq datasets. The datasets, GSE234933, GSE182227, GSE164690, and GSE181919, respectively, were selected from the Gene Expression Omnibus (GEO) platform based on having appropriated clinical annotations; these include sex, HPV status, and the origin of the sample (normal tissue, primary tumor, or metastasis)^2,16,25,32^. Prior to integration, a critical pruning process was applied to these datasets, which involved removing data points with missing essential annotations (sex, HPV status, or sample origin) and renaming variables to standardize terminology across studies. This ensured consistency and alignment in data structure, enabling more robust comparisons, and a solid foundation for accurate integration of the datasets. The final integrated dataset consists of 79 patients with complete sex and HPV status annotations, including 46 HPV-cases (32 male, 14 female) and 32 HPV+ cases (30 male, 2 female). All measurements derive from distinct biological samples across patients; no repeated measures were taken.

### Seurat Pipeline Normalization

A standard workflow using the R package `Seurat` was performed on the combined HNSCC datasets^33^. The Seurat functions help to adjust for differences in sequencing depth and technological biases between datasets. Normalization ensures that the data across different studies are compatible and can be integrated meaningfully, providing a reliable basis for comparison and further analysis. First, data normalization was performed using Seurat’s function `NormalizeData()`, which adjusts the expression measurements for each cell to allow for more accurate comparisons across different cells. Following normalization, the `FindVariableFeatures()` function was applied in order to identify the most variable genes across the dataset, which are crucial to distinguishing between cell types. Data were scaled with `ScaleData()` to ensure that highly variable genes do not overshadow the influence of genes with smaller variability.

### Stacas for Semi-supervised Data Integration

The R package `Stacas`, a semi-supervised method, was used to integrate single-cell datasets from multiple HNSCC studies^34^. This approach employs semi-supervised labeling to align datasets while preserving biologically relevant differences, such as unique cell type distributions or expression patterns specific to each dataset. `Run.STACAS()` was applied to the preprocessed datasets to identify shared anchors, representing mutual biological similarities across cells. These anchors were then used to project cells from different datasets into a shared space, minimizing batch effects and maintaining key biological features. By leveraging both labeled and unlabeled data through the `cell.labels` parameter, Stacas improves accuracy by matching similar cell types across datasets. This method standardizes data from diverse sources while preserving biological integrity, enabling a cohesive and reliable analysis.

### Automated Annotation of Classical Cell Types

The R package `scGate,` an annotation tool, was employed, which automates the identification of cell types in the datasets using predefined gating models (GMs)^35^. These GMs are based on commonly used markers for immune cells in humans and mice, such as T cells, B cells, NK cells, and myeloid populations. The models are refined with data from five annotated datasets from studies on blood or tumor samples, which are already integrated into the package^36–40^. This step ensured consistent and precise cell type classification across different datasets, essential for analyzing complex interactions within the TME.

### Tumor Cell Annotation Using the Ikarus Method

The Ikarus method, a fully supervised classifier designed to detect tumor cells with high specificity was used to accurately annotate tumor cells^17^. Ikarus applies pre-trained models on labeled gene signatures characteristic of tumor profiles, enabling precise tumor cell identification in complex cellular environments. The integrated Seurat object was prepared for Ikarus compatibility by converting it to the H5AD format with the `DietSeurat()` and `zellkonverter::SCE2AnnData` functions. By using Ikarus in Python, tumor cells were identified across the selected HNSCC datasets and the Ikarus-generated labels were integrated with the existing tumor cell annotations to enhance specificity and ensure reliable tumor cell identification.

### CNV Analysis

To compare chromosomal copy number variation (CNV) profiles between epithelial cells, originally annotated tumor cells, and cells reclassified from epithelial to tumor, inferCNV R package was used^41^. A subset of 4,818 cells—the limiting size of the relabeled epithelial group— was selected from each category for balanced analysis. Random sampling ensured equal representation across groups. The RNA count matrix was converted to a standard format, and cells were annotated into three categories: epithelial, tumor, and epithelial_tumor. The “Epithelial” group was used as the reference for CNV detection. Gene order was defined using hg38 coordinates, and a cutoff of 0.1 was applied to filter low-expression genes. Denoising and a Hidden Markov Model were used to infer CNV states. Analyses were performed on a high memory computational cluster.

### Subclassification of Immune Cells

A detailed subclassification of immune cells was performed utilizing the `ProjectTILS` framework, which is encompassed within the R package `scGate`. It employs preestablished reference labeling for critical immune cell types such as dendritic cells, CD4, and CD8 T cells.

### Co-variation Network Analysis Using CoVarNet

The CoVarNet framework, a computational approach designed to identify higher-order coordination among cell types, was applied to characterize TME organization across HNSCC samples. CoVarNet quantifies the covariance in cell type proportions across patients to infer cellular co-variation modules (CMs), reflecting coordinated shifts in cell population abundances. For this analysis, the integrated single-cell atlas was used to compute per-patient cell type proportions, which were then provided as input to CoVarNet in R. The framework constructs a covariance matrix based on these proportions and performs hierarchical clustering to define modules of co-varying cell types. Each module was subsequently weighted by its relative contribution of cell types, and module abundance scores were computed for each patient.

## Data Availability

The unified HNSCC atlas is available through the `HNSCatlas` R package, which can be accessed at https://github.com/DKFZ-E220/HNSCatlas. This package allows direct download and loading of the curated atlas (as an R object) either into the R environment or into a specified directory. The dataset is also available at the Helmholtz HIFIS repository: https://hifis-storage.desy.de:2880/Helmholtz/E220-Radioonc_biol-DKFZ/HNSCC_Atlas.rds. The complete code used for the creation of the atlas, including pre-processing, integration, and downstream analyses, is included in the inst/extdata folder of the GitHub repository to ensure full reproducibility.

## Ethical Compliance

This study reanalyzed publicly available de-identified scRNA-seq datasets with appropriate ethical approvals from the original studies (GSE234933, GSE182227, GSE164690, and GSE181919). No new human subjects were recruited, and no identifiable patient information was used.

