## Supplementary material for "A Unified Single-Cell Atlas of HNSCC: Uncovering HPV and Sex Variability in the Tumor Microenvironment": suppl_figures_tables

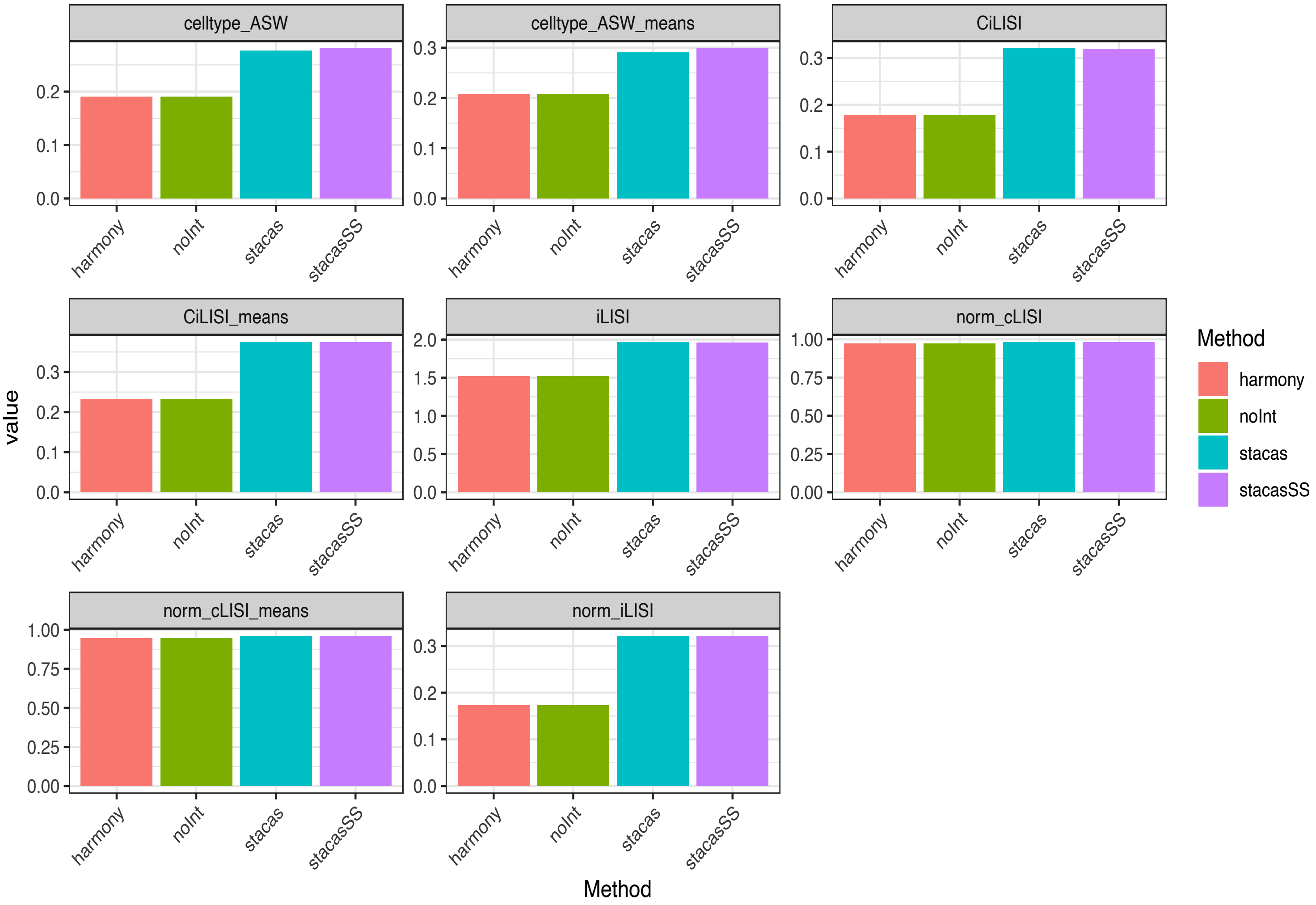


**Supplementary Figure 1. Evaluation of integration quality across methods using batch mixing and cell identity preservation metrics.**

This figure compares the performance of different data integration strategies, Harmony, no integration (noInt), STACAS, and STACAS with semi-supervised anchoring (STACAS-SS), based on a range of established metrics. These include cell type Adjusted Silhouette Width (ASW), cell-type and integration-specific Local Inverse Simpson's Index (cLISI and iLISI), and their normalized and averaged values. STACAS and STACAS-SS consistently achieve higher iLISI and ASW scores.

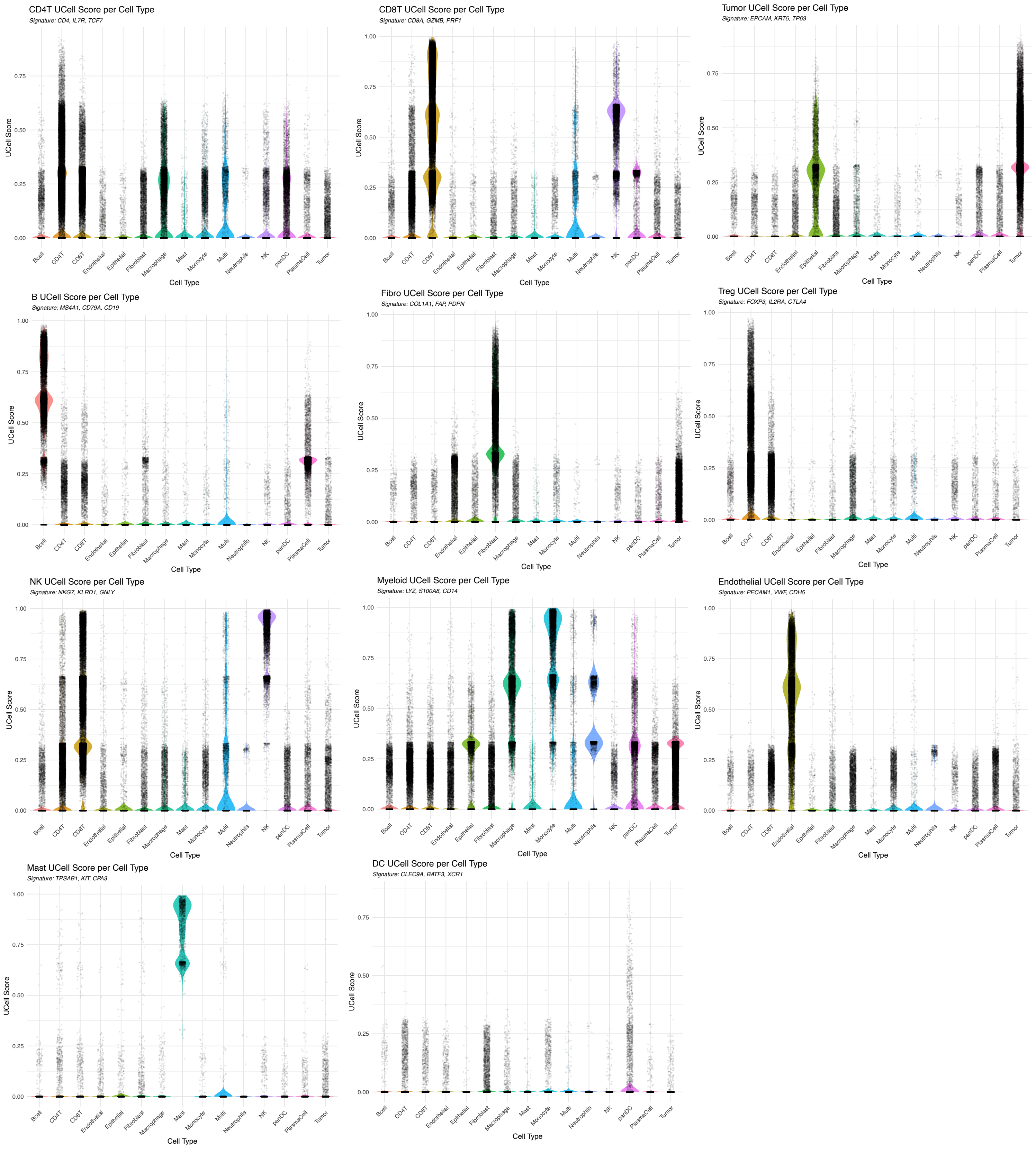


**Supplementary Figure 2. Cell type specific gene signature scoring across annotated cell populations.** Violin plots showing the expression of curated gene signatures across the final annotated cell types in the HNSCC atlas. Each panel corresponds to a distinct cell type signature (e.g., CD8⁺ T cells, CD4⁺ T cells, B cells, tumor cells or fibroblasts), and demonstrates strong enrichment of the corresponding score within the expected population.

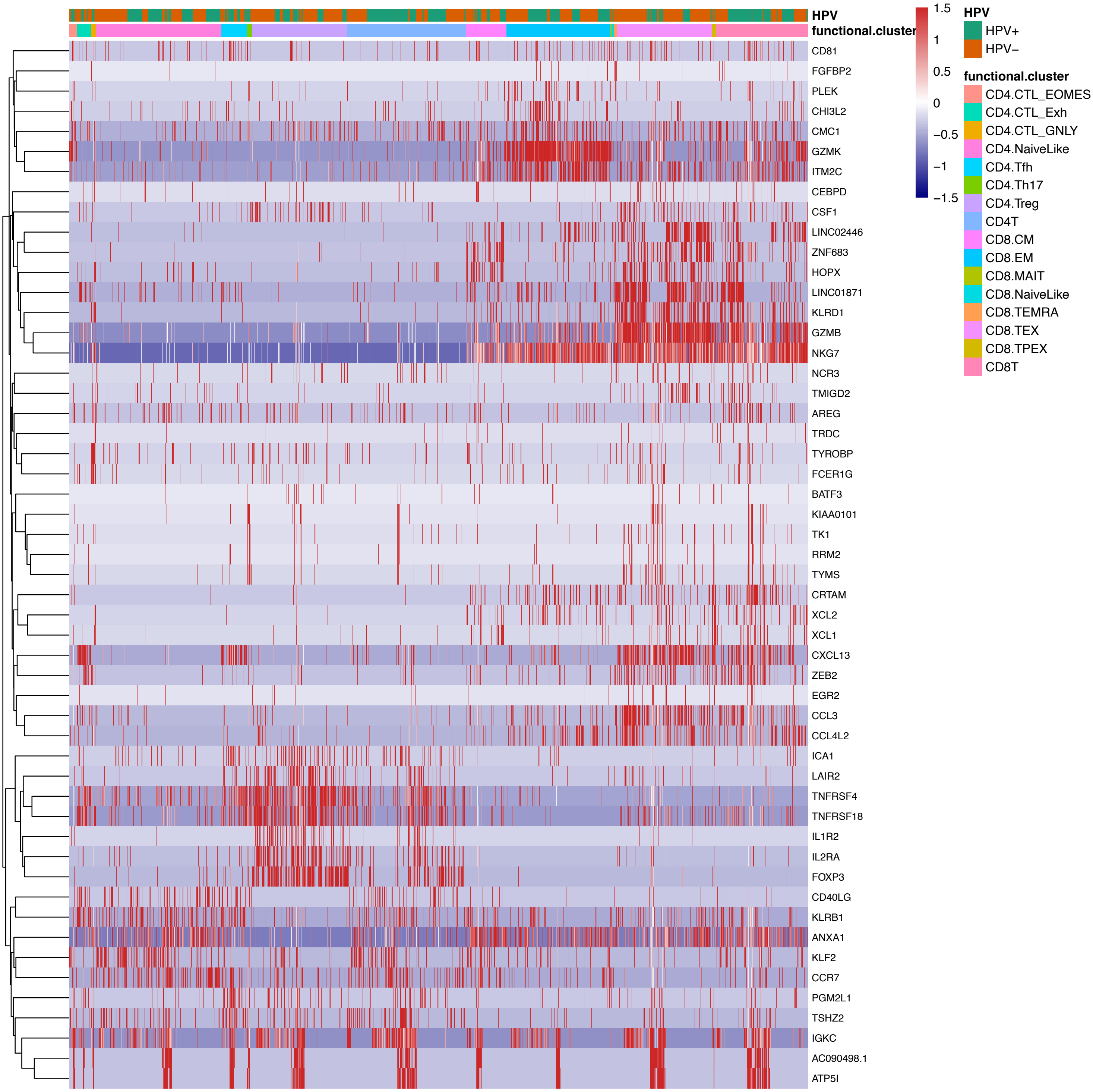


**Supplementary Figure 3. Expression of top markers across annotated T cell subtypes.**
This heatmap displays the scaled expression of the top differentially expressed genes across the annotated T cell clusters. Marker genes associated with cytotoxicity (e.g., GZMB, GZMK, NKG7), exhaustion (e.g., CXCL13, KLRD1), regulatory T cells (e.g., TNFRSF4, IL2RA, FOXP3), and helper T cell subsets (e.g., CXCL13, CD40LG, CCR7) show selective enrichment in their corresponding cell populations.

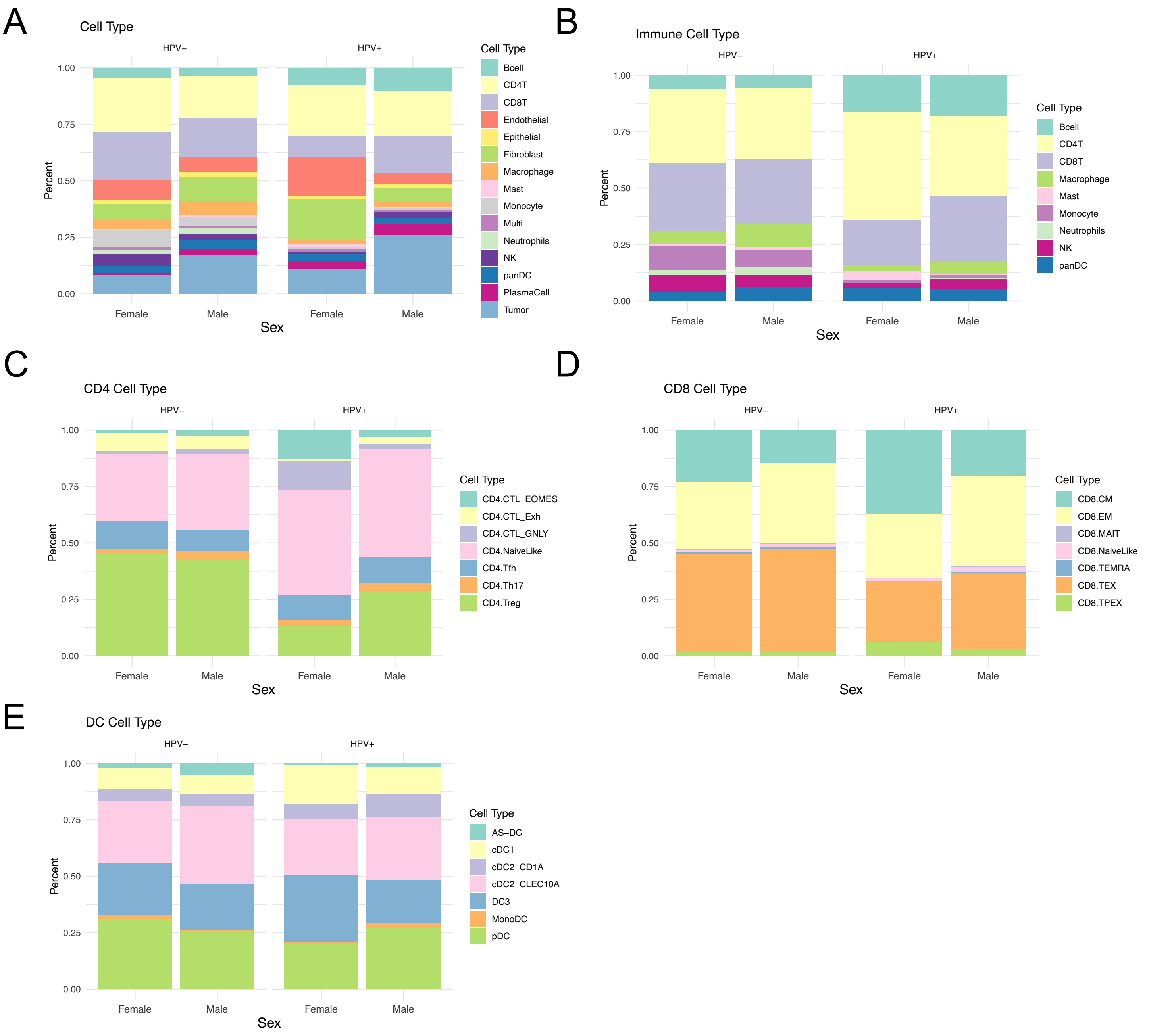


**Supplementary Figure 4. Cell type composition in HNSCC tumors by sex, stratified by HPV status.** **(A)** Stacked bar plot showing the proportion of each major cell type across female and male groups, separated by HPV status (HPV+ and HPV-). The plot highlights sex differences in immune and stromal cell distributions. **(B)** Stacked bar plot focusing on overall immune cell types, showing distribution differences between females and males in HPV+ and HPV- groups. **(C)** Distribution of CD4+ T cell subtypes across sex and HPV status. **(D)** Distribution of CD8+ T cell subtypes across sex and HPV status.**(E)** Distribution of dendritic cell (DC) subtype composition across sex and HPV status.

| Patient | Sex | HPV | Cohort | Total_cells | Origin |
|---|---|---|---|---|---|
| BHN1 | Male | HPV- | GSE234933 | 2568 | Primary |
| BHN17 | Male | HPV- | GSE234933 | 5756 | Primary |
| BHN30 | Male | HPV- | GSE234933 | 4420 | Primary |
| BHN31TS | Male | HPV- | GSE234933 | 3868 | Primary |
| BHN39 | Male | HPV- | GSE234933 | 1272 | Primary |
| BHN40-3-05 | Female | HPV- | GSE234933 | 1735 | Primary |
| BHN46 | Female | HPV- | GSE234933 | 4181 | Primary |
| BHN49 | Male | HPV- | GSE234933 | 3254 | Primary |
| BHN50 | Male | HPV+ | GSE234933 | 346 | Primary |
| BHN52 | Male | HPV+ | GSE234933 | 1921 | Primary |
| BHN58 | Male | HPV+ | GSE234933 | 2818 | Primary |
| BHN59 | Male | HPV+ | GSE234933 | 5046 | Primary |
| BHN60 | Female | HPV- | GSE234933 | 4932 | Primary |
| BHN63 | Male | HPV+ | GSE234933 | 8330 | Primary |
| BHN64 | Male | HPV- | GSE234933 | 4774 | Primary |
| BHN67 | Male | HPV- | GSE234933 | 3643 | Primary |
| BHN68 | Male | HPV+ | GSE234933 | 8450 | Primary |
| BHN7 | Male | HPV- | GSE234933 | 1081 | Primary |
| BHN70 | Male | HPV- | GSE234933 | 4739 | Primary |
| BHN72 | Male | HPV- | GSE234933 | 3962 | Primary |
| BHN74 | Female | HPV- | GSE234933 | 9194 | Primary |
| BHN75 | Male | HPV- | GSE234933 | 6981 | Primary |
| BHN76 | Male | HPV+ | GSE234933 | 6685 | Primary |
| BHN77 | Male | HPV+ | GSE234933 | 7817 | Primary |
| HN01 | Male | HPV- | GSE164690 | 3473 | Primary |
| HN02 | Female | HPV- | GSE164690 | 748 | Primary |
| HN03 | Male | HPV- | GSE164690 | 5115 | Primary |
| HN04 | Male | HPV- | GSE164690 | 1383 | Primary |
| HN05 | Female | HPV- | GSE164690 | 5111 | Primary |
| HN06 | Male | HPV- | GSE164690 | 2667 | Primary |
| HN07 | Male | HPV- | GSE164690 | 3139 | Primary |
| HN08 | Female | HPV- | GSE164690 | 2289 | Primary |
| HN09 | Female | HPV- | GSE164690 | 3701 | Primary |
| HN10 | Male | HPV- | GSE164690 | 4465 | Primary |
| HN11 | Male | HPV- | GSE164690 | 4762 | Primary |
| HN12 | Male | HPV+ | GSE164690 | 4543 | Primary |
| HN13 | Male | HPV+ | GSE164690 | 5284 | Primary |
| HN14 | Male | HPV+ | GSE164690 | 3720 | Primary |
| HN15 | Female | HPV- | GSE164690 | 4472 | Primary |
| HN16 | Male | HPV+ | GSE164690 | 5086 | Primary |
| HN17 | Male | HPV+ | GSE164690 | 10667 | Primary |
| HN18 | Male | HPV+ | GSE164690 | 2455 | Primary |
| OP10 | Male | HPV- | GSE182227 | 7078 | Primary |
| OP12 | Male | HPV- | GSE182227 | 6911 | Primary |
| OP13 | Male | HPV+ | GSE182227 | 744 | Primary |
| OP14 | Male | HPV+ | GSE182227 | 3238 | Primary |
| OP16 | Male | HPV- | GSE182227 | 3330 | Primary |
| OP17 | Male | HPV+ | GSE182227 | 1163 | Primary |
| OP19 | Female | HPV- | GSE182227 | 2274 | Primary |
| OP20 | Male | HPV+ | GSE182227 | 6187 | Primary |
| OP33 | Male | HPV+ | GSE182227 | 7212 | Primary |
| OP34 | Male | HPV+ | GSE182227 | 7232 | Primary |
| OP35 | Male | HPV+ | GSE182227 | 1167 | Primary |
| OP4 | Male | HPV+ | GSE182227 | 5006 | Primary |
| OP5 | Male | HPV+ | GSE182227 | 2664 | Primary |
| OP6 | Female | HPV+ | GSE182227 | 3269 | Primary |
| OP8 | Male | HPV- | GSE182227 | 6964 | Primary |
| OP9 | Male | HPV+ | GSE182227 | 6531 | Primary |
| P15 | Female | HPV- | GSE181919 | 1731 | Primary |
| P21 | Male | HPV- | GSE181919 | 697 | Primary |
| P22 | Male | HPV+ | GSE181919 | 1143 | Primary |
| P26 | Male | HPV- | GSE181919 | 944 | Primary |
| P30 | Male | HPV- | GSE181919 | 282 | Primary |
| P31 | Male | HPV- | GSE181919 | 523 | Primary |
| P38 | Male | HPV- | GSE181919 | 1049 | Primary |
| P4 | Female | HPV- | GSE181919 | 608 | Primary |
| P43 | Male | HPV+ | GSE181919 | 1662 | Primary |
| P46 | Male | HPV+ | GSE181919 | 1413 | Primary |
| P51 | Male | HPV- | GSE181919 | 1801 | Primary |
| P57 | Male | HPV+ | GSE181919 | 1104 | Primary |
| P59 | Male | HPV+ | GSE181919 | 2350 | Primary |
| P6 | Male | HPV- | GSE181919 | 1327 | Primary |
| P60 | Female | HPV- | GSE181919 | 1948 | Primary |
| P7 | Male | HPV- | GSE181919 | 536 | Primary |
| P8 | Male | HPV- | GSE181919 | 1206 | Primary |
| P84 | Male | HPV+ | GSE181919 | 830 | Primary |
| P86 | Female | HPV+ | GSE181919 | 1084 | Primary |
| P9 | Female | HPV- | GSE181919 | 850 | Primary |

**Supplementary Table 1. Clinical and technical characteristics of HNSCC patient samples included in the single-cell atlas.** The table includes information on patient sex, HPV status, dataset of origin, total number of single cells retained after filtering, and tissue source (e.g., primary tumor, lymph node, or other anatomical location).

2

**Percentage All Cell Types**

| | HPV - | | HPV + | |
|---|---|---|---|---|
| | Female | Male | Female | Male |
| Bcell | 4.4% | 3.6% | 7.7% | 10.20% |
| CD4T | 23.9% | 18.7% | 22.5% | 20.00% |
| CD8T | 21.7% | 17.2% | 9.5% | 16.30% |
| Endothelial | 8.9% | 6.8% | 17.0% | 4.90% |
| Epithelial | 1.6% | 2.0% | 1.6% | 2.00% |
| Fibroblast | 6.7% | 10.9% | 18.3% | 5.50% |
| Macrophage | 4.2% | 5.9% | 1.3% | 2.90% |
| Mast | 0.5% | 0.9% | 1.7% | 0.40% |
| Monocyte | 7.8% | 4.2% | 0.8% | 0.80% |
| Multi | 1.2% | 0.9% | 1.4% | 0.90% |
| NK | 5.5% | 3.1% | 1.0% | 2.70% |
| Neutrophils | 1.6% | 2.2% | 0.0% | 0.00% |
| PlasmaCell | 0.9% | 3.0% | 3.5% | 4.60% |
| Tumor | 8.4% | 16.9% | 11.2% | 26.10% |
| panDC | 2.9% | 3.7% | 2.7% | 2.90% |

3

**Percentage Immune Cell Types**

| | HPV - | | HPV + | |
|---|---|---|---|---|
| | Female | Male | Female | Male |
| Bcell | 6.0% | 6.0% | 16.3% | 18.2% |
| CD4T | 33.0% | 31.4% | 47.7% | 35.6% |
| CD8T | 29.9% | 28.8% | 20.1% | 29.0% |
| Macrophage | 5.9% | 9.9% | 2.8% | 5.2% |
| Mast | 0.7% | 1.5% | 3.7% | 0.7% |
| Monocyte | 10.7% | 7.1% | 1.6% | 1.4% |
| NK | 7.5% | 5.2% | 2.1% | 4.8% |
| Neutrophils | 2.3% | 3.8% | 0.0% | 0.1% |
| panDC | 4.0% | 6.3% | 5.7% | 5.1% |

4

**Percentage CD8 Cell Types**

| | HPV - | | HPV + | |
|---|---|---|---|---|
| | Female | Male | Female | Male |
| CD8.CM | 23.0% | 14.8% | 37.1% | 20.3% |
| CD8.EM | 29.6% | 35.4% | 28.5% | 40.2% |
| CD8.MAIT | 0.5% | 0.4% | 0.0% | 0.4% |
| CD8.NaiveLike | 0.8% | 1.0% | 1.3% | 2.2% |
| CD8.TEMRA | 1.2% | 1.3% | 0.3% | 0.4% |
| CD8.TEX | 43.3% | 45.5% | 26.6% | 33.6% |
| CD8.TPEX | 1.6% | 1.6% | 6.2% | 3.0% |

5

**Percentage CD4 Cell Types**

| | HPV - | | HPV + | |
|---|---|---|---|---|
| | Female | Male | Female | Male |
| CD4.CTL_EOMES | 1.2% | 2.7% | 12.8% | 2.9% |
| CD4.CTL_Exh | 8.0% | 6.0% | 1.2% | 3.5% |
| CD4.CTL_GNLY | 1.5% | 2.0% | 12.5% | 2.1% |
| CD4.NaiveLike | 29.5% | 33.8% | 46.3% | 47.9% |
| CD4.Tfh | 12.4% | 9.3% | 11.2% | 11.5% |
| CD4.Th17 | 2.0% | 3.9% | 2.8% | 3.2% |
| CD4.Treg | 45.5% | 42.2% | 13.1% | 28.9% |

6

**Percentage DC Cell Types**

| | HPV - | | HPV + | |
|---|---|---|---|---|
| | Female | Male | Female | Male |
| AS-DC | 2.3% | 5.0% | 1.0% | 1.5% |
| DC3 | 23.1% | 20.5% | 29.5% | 18.9% |
| MonoDC | 1.7% | 0.9% | 1.0% | 2.4% |
| cDC1 | 9.2% | 8.4% | 17.1% | 12.1% |
| cDC2_CD1A | 5.4% | 5.7% | 6.7% | 10.1% |
| cDC2_CLEC10A | 27.4% | 34.6% | 24.8% | 28.1% |
| pDC | 30.9% | 25.0% | 20.0% | 27.0% |

**Supplementary Tables 2-6. Cell type composition in HNSCC tumors by sex, stratified by HPV status.**
(2) Table showing the proportion of each major cell type across female and male groups, separated by HPV status (HPV+ and HPV−). (3) Table showing the proportion of each major immune cell type across female and male groups, separated by HPV status (4) Table showing the proportion of $CD8^{+}$ T cell subtypes across female and male groups, separated by HPV status (5) Table showing the proportion of $CD4^{+}$ T cell subtypes across female and male groups, separated by HPV status (6) Table showing the proportion of DC subtypes across female and male groups, separated by HPV status

| | Cell Type | Description |
|---|---|---|
| 6 | CD8.NaiveLike | Antigen-naive T cells |
| | CD8.CM | Central Memory T cells |
| | CD8.EM | Effector Memory T cells |
| | CD8.TEMRA | Effector Memory cells re-expressing CD45RA. Sometimes called Short Lived Effectors (SLEC), or Cytotoxic effectors |
| | CD8.TPEX | Progenitor exhausted T cells |
| | CD8.TEX | Exhausted T cells |
| | CD8.MAIT | Mucosal-associated invariant T cells, innate-like T cells defined by their semi-invariant αβ T cell receptor |
| | CD4.NaiveLike | T cells with naive-like phenotype |
| | CD4.Tfh | T follicular helper cells |
| | CD4.Th17 | Th17 helper cells |
| | CD4.Treg | T regulatory cells |
| | CD4.CTL_EOMES | Cytotoxic CD4 T cells expressing EOMES and GZMK |
| | CD4.CTL_GNLY | Cytotoxic CD4 T cells expressing GNLY |
| | CD4.CTL_Exh | Cytotoxic CD4 T cells with exhaustion phenotype |
| | AS-DC | AXL+ SIGLEC6+ Dendritic Cells, also referred to as DC5 or Pre-DCs |
| | cDC1 | Conventional Dendritic Cells type 1, specialized in antigen cross-presentation and CD8+ T cell activation |
| | cDC2_CD1A | Conventional Dendritic Cells type 2 subset expressing CD1A, involved in CD4+ T cell activation |
| | cDC2_CLEC10A | Conventional Dendritic Cells type 2 subset expressing CLEC10A, functionally distinct from CD1A-expressing cDC2 |
| | DC3 | Tissue-resident DCs, lacking a direct counterpart in circulation, potentially derived from cDC1 or cDC2/MoDC |
| | MonoDC | Monocyte-derived Dendritic Cells, inflammation-induced, with similarities to cDC2 |
| | pDC | Plasmacytoid Dendritic Cells, key producers of type I IFNs, fostering antitumor immunity |

**Supplementary Table 7. Overview of $CD8^+$, $CD4^+$, and dendritic cell subtypes.**

This table provides an overview of major $CD8^+$ T cell, $CD4^+$ T cell, and dendritic cell (DC) subtypes, including their definitions and functional roles in immune regulation. This table serves as a reference for the classification and functional annotation of immune cell subsets in HNSCC tumors, facilitating comparisons across different studies.
